# Ecogenomics and biogeochemical impacts of uncultivated globally abundant ocean viruses

**DOI:** 10.1101/053090

**Authors:** Simon Roux, Jennifer R. Brum, Bas E. Dutilh, Shinichi Sunagawa, Melissa B. Duhaime, Alexander Loy, Bonnie T. Poulos, Natalie Solonenko, Elena Lara, Julie Poulain, Stéphane Pesant, Stefanie Kandels-Lewis, Céline Dimier, Marc Picheral, Sarah Searson, Corinne Cruaud, Adriana Alberti, Carlos M. Duarte, Josep M. Gasol, Dolors Vaqué, Tara Oceans Coordinators, Peer Bork, Silvia G. Acinas, Patrick Wincker, Matthew B. Sullivan, Silvia G. Acinas, Peer Bork, Emmanuel Boss, Chris Bowler, Colomban de Vargas, Michael Follows, Gabriel Gorsky, Nigel Grimsley, Pascal Hingamp, Daniele Iudicone, Olivier Jaillon, Stefanie Kandels-Lewis, Lee Karp-Boss, Eric Karsenti, Uros Krzic, Fabrice Not, Hiroyuki Ogata, Stephane Pesant, Jeroen Raes, Emmanuel G. Reynaud, Christian Sardet, Mike Sieracki, Sabrina Speich, Lars Stemmann, Matthew B. Sullivan, Shinichi Sunagawa, Didier Velayoudon, Patrick Wincker

**Author notes:** Tara* Oceans coordinators and affiliations are listed following the Acknowledgements. Current address: National Science Foundation, Arlington, Virginia, USA. Current address: Department of Geosciences, Laboratoire de Météorologie Dynamique (LMD), Ecole Normale Supérieure, 24 rue Lhomond 75231 Paris, Cedex 05, France.

## Abstract

Ocean microbes drive global-scale biogeochemical cycling^1^, but do so under constraints imposed by viruses on host community composition, metabolism, and evolutionary trajectories^2–5^. Due to sampling and cultivation challenges, genome-level viral diversity remains poorly described and grossly understudied in nature such that <1% of observed surface ocean viruses, even those that are abundant and ubiquitous, are ‘known’^5^. Here we analyze a global map of abundant, double stranded DNA (dsDNA) viruses and viral-encoded auxiliary metabolic genes (AMGs) with genomic and ecological contexts through the Global Ocean Viromes (GOV) dataset, which includes complete genomes and large genomic fragments from both surface and deep ocean viruses sampled during the *Tara* Oceans and *Malaspina* research expeditions^6,7^. A total of 15,222 epi- and mesopelagic viral populations were identified that comprised 867 viral clusters (VCs, approximately genus-level groups^8,9^). This roughly triples known ocean viral populations^10^, doubles known candidate bacterial and archaeal virus genera^9^, and near-completely samples epipelagic communities at both the population and VC level. Thirty-eight of the 867 VCs were identified as the most impactful dsDNA viral groups in the oceans, as these were locally or globally abundant and accounted together for nearly half of the viral populations in any GOV sample. Most of these were predicted *in silico* to infect dominant, ecologically relevant microbes, while two thirds of them represent newly described viruses that lacked any cultivated representative. Beyond these taxon-specific ecological observations, we identified 243 viral-encoded AMGs in GOV, only 95 of which were known. Deeper analyses of 4 of these AMGs revealed that abundant viruses directly manipulate sulfur and nitrogen cycling, and do so throughout the epipelagic ocean. Together these data provide a critically-needed organismal catalog and functional context to begin meaningfully integrating viruses into ecosystem models as key players in nutrient cycling and trophic networks.

---

The fundamental bottleneck preventing the incorporation of viruses of microbes into predictive ecosystem models is the lack of quantitative surveys of viral diversity in nature. This is because (i) most naturally-occurring microbes and viruses resist being cultured, and (ii) viruses lack a universally conserved marker gene, which precludes barcode surveys of uncultivated viral diversity^5^. While viral metagenomics was introduced to circumvent these issues, low-throughput sequencing technologies initially yielded highly fragmented datasets suitable only for strongly database-biased descriptions^11^, and gene-level analyses of environmental viral communities (reviewed in ref. 5).

Subsequent improvements in experimental methods, sequencing technologies, and analytical approaches progressively enabled viral population ecology through the availability of genomic information^5,12–14^. For example, 1,148 large viral genome fragments captured in a fosmid library of Mediterranean Sea microbes revealed remarkable viral diversity in a single sample, with some of these genomes seemingly globally distributed based upon the six viral metagenomic datasets available at the time^12^. Similarly, 69 viral genomes assembled from single-cell genomic datasets provided reference genomes that illuminated the ecology, evolution and biogeochemical impacts of viruses infecting an uncultivated anaerobic chemoautotroph^14^. Beyond these ‘omics-enabled experimental advances, metagenomic approaches have matured to be quantitative^5^ and informative enough, at least for dsDNA templates, to themselves provide genomic information on viruses that infect both abundant and rare hosts. For example, the analysis of 43 surface ocean viral metagenomes (viromes) comprising the *Tara* Oceans Viromes (TOV) dataset revealed the global underlying structure of these communities, and identified 5,476 viral populations, only 39 of which were previously known^10^.

Here we further identify ocean viral populations, characterize the most abundant and widespread types of ocean viruses, and analyze new viral-encoded AMGs and their distributions to expand our understanding of how viruses modulate microbial biogeochemistry. We do so on the basis of a new Global Oceans Viromes (GOV) dataset, which augments TOV with 61 new samples to better represent the surface and deep oceans, and now totals 104 ocean viromes representing 925 Gbp of sequencing (Supplementary Table 1). Beyond better sample coverage, analytical approaches including cross-assembly^15^ and genome binning^16^ make the GOV dataset a much improved genomic representation of the sampled viruses. From 1,380,834 contigs generated, which recruited 67% of the reads, we identified 15,280 viral populations (Fig. 1A, Supplementary Fig. 1). This expands the number of known ocean viral populations nearly 3-fold over the prior TOV dataset^10^, while also improving the genomic context for these TOV-known populations by a 2.5-fold increase in contig length on average (Supplementary Table 2). Rarefaction analyses show that while mesopelagic viral communities remain undersampled, epipelagic viral communities now appear near-completely sampled (Extended Data Fig. 1A). Because bathypelagic communities were underrepresented due to cellular contamination of these viromes, we focused the remaining analyses on the 15,222 non-bathypelagic viral populations.

**Figure 1:**
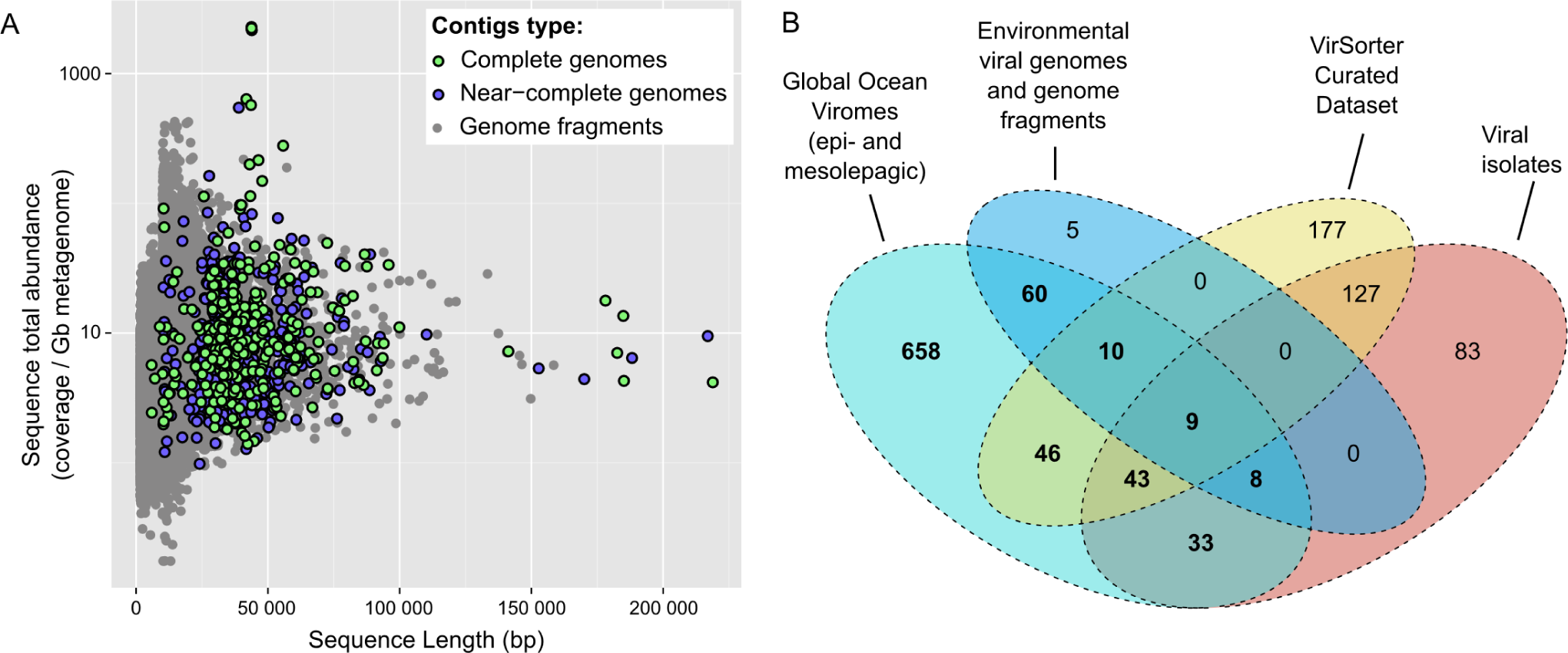
Composition and novelty of the Global Ocean Viromes (GOV) dataset. **A.** Size of viral contigs (x-axis) and cumulative coverage across the GOV dataset (y-axis). Contigs corresponding to complete or near-complete genomes (based on the size of similar complete genomes) are indicated. For clarity, only contigs associated with a viral population are displayed. **B.** Distribution of all viral clusters (VCs) according to the origin of their members. Viral genomes (or fragments) in a VC can originate from isolate viral genomes, the VirSorter Curated Dataset^9^ (viral genomes identified *in silico* from microbial genomes), environmental viral genomes and genome fragments (e.g. from fosmid libraries), or the GOV dataset. VCs including at least one GOV sequence and further analyzed in this study are highlighted in bold.

We first categorized new and known viral populations into viral clusters, or VCs (Supplementary Fig. 1) using shared gene content information and network analytics^8^. This method starts from genome fragments (≥10kb) and results in VCs approximately equivalent to known bacterial and archaeal virus genera^8,9^ (see also Supplementary Text, Extended Data Fig. 2 & Supplementary Table 3 for comparison with alternative classification methods). Combining the 15,222 viral populations identified here with the genomes and genome fragments of another 15,929 publicly available bacterial and archaeal viruses generated a total of 1,259 VCs. Of these, 658 included exclusively GOV sequences, which approximately doubles known bacterial and archaeal virus genera^9^, and another 209 VCs contained at least one GOV sequence (Fig. 1B). As with viral populations, rarefaction analyses suggested that VC diversity was undersampled in mesopelagic waters, but near-completely sampled in epipelagic waters (Extended Data Fig. 1B).

We next identified the most abundant and widespread VCs based on read recruitment of VC members. In each sample, a fraction of the VCs were identified as abundant based on their cumulative contribution to sample diversity (estimated with the Simpson Index, abundant VCs represent 80% of the total sample diversity, Extended Data Fig. 1C). By these criteria, only 38 of the 867 VCs observed were abundant in two or more stations, and together recruited an average of 50% and 35% of the reads from viral populations for epi- and mesopelagic samples, respectively (Supplementary Table 3). Four of these 38 abundant VCs were also relatively ubiquitous as they were abundant in more than 25 stations, and 62 of the 91 non-bathypelagic samples were dominated by 1 of these 4 VCs (Fig. 2 A & B). Among the 38 abundant VCs, only 2 corresponded to well-studied viruses, from the T4 superfamily^17,18^ (VC_2, 1 of the 4 ubiquitous) and the *T7virus* genus^19^ (VC_9), whereas 8 represented known, but unclassified viral isolates, another 10 included viruses previously only identified in environmental libraries^12,13^, and the remaining 18 VCs were completely novel (Fig. 2C, Extended Data Fig. 3).

**Figure 2:**
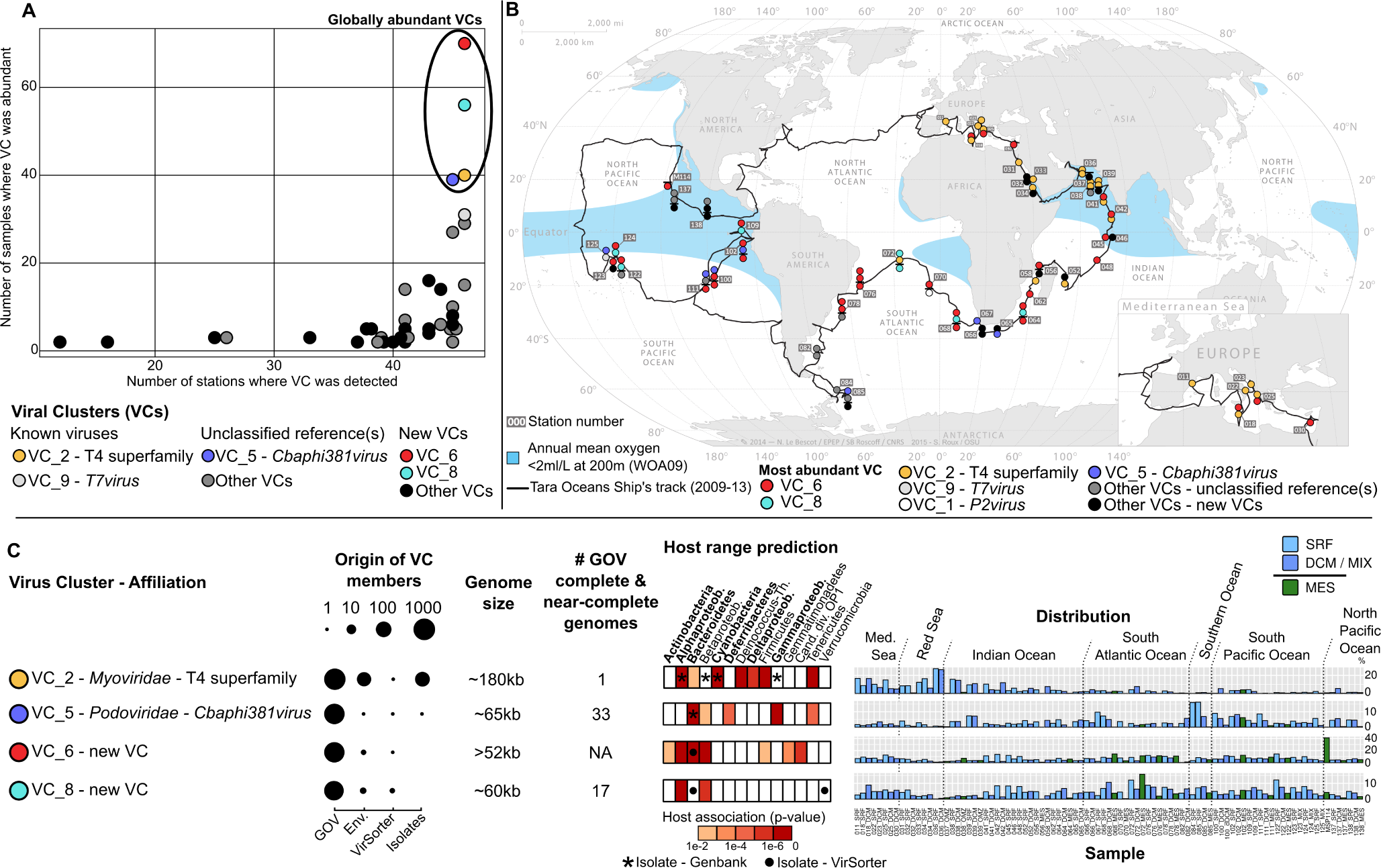
Characterization of the dominant oceanic viral clusters (VCs) **A.** Distribution and abundance of the 38 recurrently abundant VCs according to the total number of stations in which members of the VC were detected (x-axis), and the number of samples in which the VC was detected in the abundant fraction (y-axis). VCs are classified by degree of novelty: “Known viruses” are VCs corresponding to established viral groups in ICTV classification, “Unclassified reference(s)” are VCs including genomes from unclassified isolate(s), and “New VCs” are only composed of environmental viral sequences (i.e. no isolate). The 4 widespread and abundant VCs are highlighted with colored circles. **B.** GOV samples with their most abundant VC mapped to station locations. Samples are stacked vertically when multiple samples from different depths are available (a horizontal line is used to separate epipelagic from mesopelagic samples). **C.** Summary of VC affiliation, origin of the VC members (Env: environmental viral sequences), estimated genome size, predicted host range, and distribution of the 4 ubiquitous VCs circled in panel A (relative abundance are indicated as % of the viral populations identified). The abundant epipelagic microbial groups (representing >1% of the microbial OTUs abundance of epipelagic samples) are highlighted in bold; Alphaproteob.-Alphaproteobacteria, Betaproteob.-Betaproteobacteria, Deinococcus-Th.-Deinococcus-Thermus, Deltaproteob.-Deltaproteobacteria, Gammaproteob.-Gammaproteobacteria, Cand div OP1-Candidate division OP1. Oceanic basins are indicated for VCs distributions; Med. Sea-Mediterranean Sea.

Given this global map of the dominant dsDNA viral types in the oceans, we next sought to identify the range of hosts that these viruses infect. Large-scale host range estimations are challenging as culture-based methods experimentally link viruses and hosts, but insufficiently capture naturally-occurring diversity, whereas metagenomic approaches broadly survey diversity, but struggle to establish virus-host linkages. However, recent work has demonstrated the predictive power of sequence-based approaches such as similarities between (i) viral genomes and host CRISPR spacers^20^ (ii) viral and microbial genomes due to integrated prophages or gene transfers^12^ and (iii) viral and host genome nucleotide signatures (here, tetranucleotide frequencies)^9^. We applied all 3 methods to the GOV dataset and predicted hosts at the phylum level, or class level for Proteobacteria (Supplementary Table 4). These results were then summarized at the VC level, leading to host range predictions for 392 of 867 VCs, and for which confidence was assessed by comparison to a null model (Supplementary Fig. 1).

The hosts of the 38 globally abundant VCs were largely restricted to abundant and widespread epipelagic-ocean microbes identified from _mi_Tag-based OTU counts in *Tara* Oceans microbial metagenomes^21^. Notably, the 4 ubiquitous and abundant VCs were predicted to infect 7 of the 8 globally abundant microbial groups (Actinobacteria, Alpha-, Delta-, and Gamma-proteobacteria, Bacteroidetes, Cyanobacteria, Deferribacteres; Fig. 2C, Extended Data Fig. 4). The 8[th] abundant microbial group, Euryarchaeota, was not linked to these 4 VCs, but was predicted as a host for 2 of the 34 other abundant VCs (VC_3 and VC_63, Extended Data Fig. 3). Among the 38 abundant VCs, the number of VCs predicted to infect a given microbial host phylum (or class for Proteobacteria) was positively correlated with host global richness rather than abundance (Extended Data Fig. 4B). This suggests that, likely because of the global distribution of ocean viruses^10,22^, widespread and abundant hosts that are minimally diverse (e.g. Cyanobacteria) provide few viral niches, whereas more diverse host groups, even at lower abundance (e.g. Betaproteobacteria), provide more opportunity for viral niche differentiation. These data thus provide critically-needed empirical support for guiding and testing hypotheses derived from global virus-host network models^23^.

Having mapped viral diversity and predicted virus-host pairings, we next sought to identify novel virus-encoded auxiliary metabolic genes, or AMGs, that might modify host metabolism during infection. To maximize the detection of AMG, all viral contigs >1.5kb were examined, including small contigs not associated with a viral population, which totaled 298,383 contigs. This revealed 243 putative AMGs (Supplementary Table 5). While 95 were already known (reviewed in ref. 24), others offer new insights into how viruses directly manipulate microbial metabolisms beyond photosynthesis and carbon metabolism^25–28^. Here we focus specifically on 4 AMGs (*dsr*C, *sox*YZ, P-II and *amo*C, see Extended Data Table 1) because of their critical roles in sulfur or nitrogen cycling and their novelty in epipelagic ocean viruses. Three of these are not yet known in viruses, and one, *dsrC*-like genes, has been observed in viruses, but only from anoxic deep-sea environments^14,29^.

Sulfur oxidation in seawater involves two central microbial pathways – Dsr and Sox^30^ – and GOV AMG analyses revealed that epipelagic viruses encode key genes for each. First, 11 *dsrC-*like genes were identified in viral contigs (Extended Data Fig. 5). The Dsr operon is used by sulfate/sulfite-reducing microbes in anoxic environments, as well as sulfur-oxidizing bacteria in oxic and anoxic environments (Fig. 3A)^30,31^. DsrC, specifically, dictates sulfur metabolism rates, as it provides the sulfur substrate to DsrAB-sulfite reductase for processing^32^. A conserved C-terminal motif with two cysteine residues Cys_B_X_10_Cys_A_ is essential for this function. Outside of energy metabolism, a second class of *dsr*C-like genes (also known as *tus*E) lack Cys_B_ and are instead involved in tRNA modification^33^. In the GOV dataset, four clades of *dsrC*-like sequences were similar to TusE (DsrC-1 to DsrC-4), whereas the fifth (DsrC-5) was most similar to *bona fide* DsrC involved in sulfur oxidation (Extended Data Fig. 5, Extended Data Table 1, Supplementary Fig. 2). Second, 4 *soxYZ* genes from the *sox* operon were identified on viral contigs (Extended Data Fig. 6)^30,31^. Like DsrC, SoxYZ is an important sulfur carrier protein harboring a sulfur interaction motif identified in GOV SoxYZ proteins (Fig. 3A, Supplementary Fig. 3)^34^.

**Figure 3:**
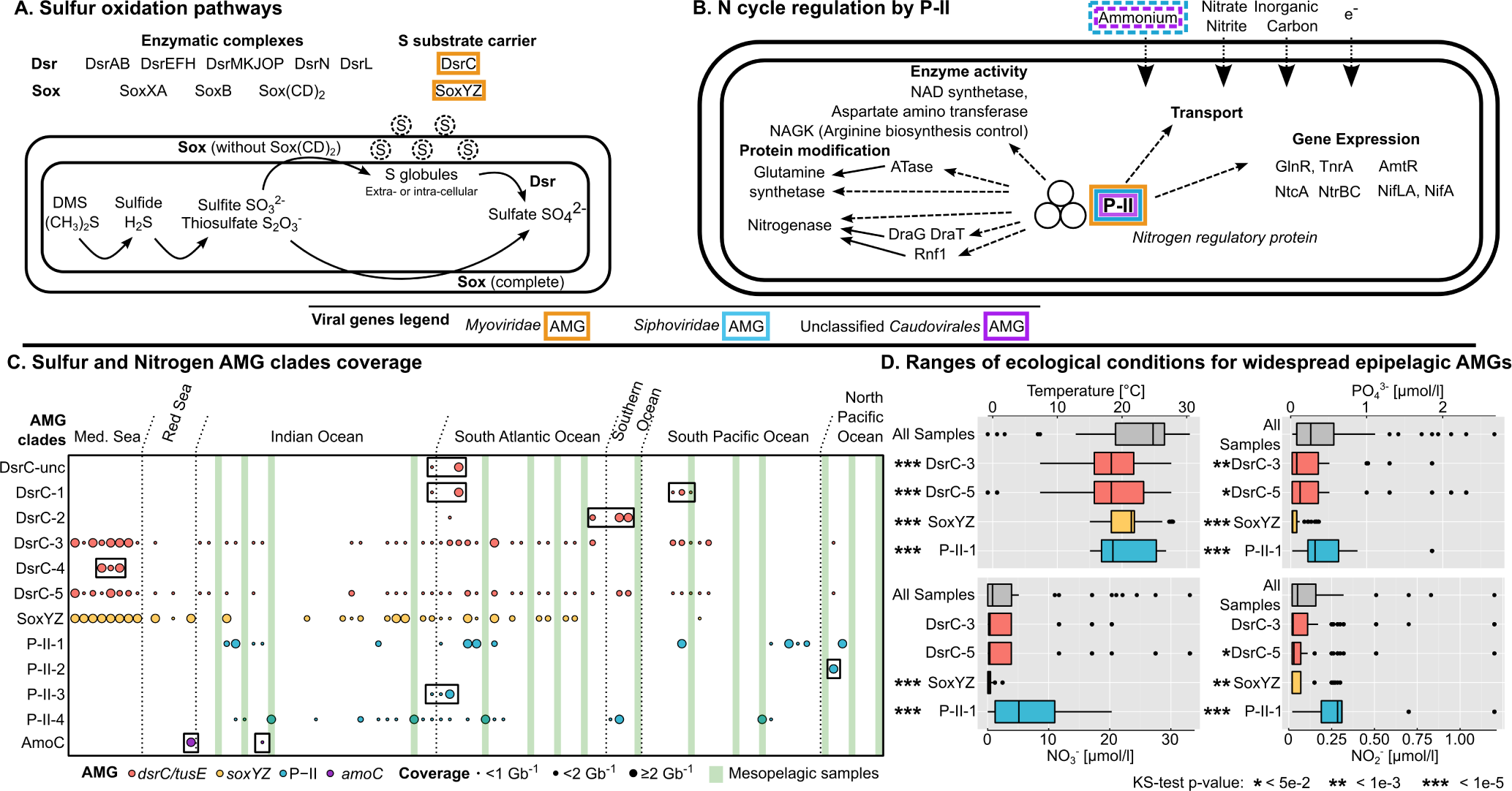
Characterization and distribution of viral Auxiliary Metabolic Genes (AMGs) involved in sulfur and nitrogen cycles. Schematics are presented for (**A**) microbial sulfur oxidation pathways involving the two main gene clusters (*dsr* and *sox*) and (**B**) the central role of the P-II protein in cell regulation (adapted from^35,45^). AMGs are outlined in colors according to the taxonomic affiliation (family) of the corresponding virus. Ammonium transporters detected next to viral P-II are highlighted with a dashed outline. **C.** Distribution of viral AMG clades. Mesopelagic samples are highlighted in dark green, and clades with restricted geographic distributions are highlighted with black boxes. **D.** Temperature and nutrient conditions for which widespread epipelagic AMGs tend to be most abundant. For each environmental parameter, the range of values across all epipelagic samples is displayed as “All Samples” alongside distributions representing the range of values where each AMG clade was detected, weighted by the AMG coverage across these samples (see Extended Data Fig. 9 for underlying coverage data). Distributions significantly different from the “All Samples” distribution (two-sided KS-test) are indicated with stars.

Complementarily, GOV AMG analyses suggested that viruses also manipulate nitrogen cycling. First, 10 GOV contigs encoded P-II, a nitrogen metabolism regulator that is widespread across bacteria and archaea (Fig. 3B)^35,36^. Functional P-II genes contain a conserved C-terminal motif, and a conserved Y residue that is uridylylated under nitrogen-limiting conditions^35^. One AMG clade (P-II-3) lacked this conversed Y residue leaving its function ambiguous, whereas 3 clades (P-II-1, P-II-2, and P-II-4) displayed both conserved motifs and were also predicted to have structures similar to bona fide P-II, which suggests these AMGs are functional (Supplementary Fig. 4). Second, two of these P-II AMG clades (P-II-1 and P-II-4) were proximal to an ammonium transporter gene, *amt*, in viral contigs (Extended Data Fig. 7). In bacteria, such an arrangement is a signature of P-II-like genes that specifically activate alternative nitrogen production and ammonia uptake pathways during nitrogen starvation^35,36^. Third, one GOV contig included *amo*C, the gene coding for the subunit C of ammonia monooxygenase, suggesting a role in ammonia oxidation^37,38^. While functional annotation is challenging for these genes^38^, and functional motifs are not yet available, the translated AMG was 94% identical to functional AmoC from Thaumarchaeota (Extended Data Fig. 8, Supplementary Fig. 5). Such exceptionally high identity is rarely observed among AMGs, and compars only to the well-studied PsbA genes, which are expressed and functional^39^. We posit that this highly conserved *amo*C AMG is also functional.

Next, we investigated the origin, evolutionary history, and diversity of these AMGs in epipelagic viruses. The 15 GOV contigs encoding *dsrC* or *soxYZ* genes were all affiliated to T4 superfamily contigs, one of the 4 abundant and ubiquitous VCs (VC_2, Extended Data Fig. 5 and 6, Extended Data Table 1). Both DsrC and Sox phylogenies suggested that these viruses obtained each AMG only once from probable S-oxidizing proteobacterial hosts (Extended Data Fig. 5 and 6). Among the latter, DsrC-5, the bona fide S-oxidation DsrC, appeared most closely related to a clade of uncultivated S-oxidizing Gammaproteobacteria represented by bacterial artificial chromosome MED13k09 (confirmed by phylogenetic analyses of the sulfur oxidizer marker DsrA, Supplementary Fig. 6). If DsrC-5-containing viruses indeed infect members of this clade, they would impact bacteria that are widespread in the epipelagic ocean^40^, and suspected to degrade dimethyl sulfide, the most important reduced sulfur species in oxygenated ocean waters and key compound in the transport of sulfur from ocean to atmosphere and in cloud formation^41^. In contrast to the sulfur AMGs, phylogenies suggest that P-II AMGs originated from diverse viruses (6 VCs including the abundant VC_2 and VC_12), and were acquired at least 4 times independently from different host phyla (Bacteroidetes, Proteobacteria, and possibly Verrucomicrobia, Extended Data Fig. 7). Finally, while a single *amoC* AMG offers only preliminary evaluation of its evolutionary history, this *amo*C-encoding contig appears to represent novel and rare unclassified archaeal dsDNA viruses (VC_623), which presumably infect ammonia-oxidizing Thaumarchaeota, known for their major role in global nitrification^37^ (Extended Data Fig. 8).

Finally, we investigated the ecology of viruses encoding these new AMGs by mapping their distribution across the GOV dataset. Seven AMG clades were geographically restricted (DsrC*-*unc, DsrC-1, DsrC-2, DsrC-4, P-II-2, P-II-3, and *amo*C), and 5 were widespread throughout the epipelagic (DsrC-3, DsrC-5, SoxYZ, P-II-1) or mesopelagic (P-II-4) ocean (Fig. 3C). While all widespread epipelagic AMGs were detected in waters of mid-range temperatures, DsrC-5 and SoxYZ were predominantly detected in low nutrients conditions (low phosphate and nitrite), while P-II-1 was predominantly detected in high nutrient conditions (high phosphate, nitrate and nitrite, Fig. 3D, Extended Data Fig. 9). Thus, we hypothesize that viruses utilize DsrC-5 or SoxYZ to boost sulfur oxidation rates when infecting sulfur oxidizers in low-nutrient conditions, and P-II under high-nutrient conditions favorable for normal host growth. The latter could be particularly useful to viruses by activating expensive and stress-inducing alternative N-producing pathways typically only used under N-starvation conditions^35,36^. Consistent with this, metatranscriptomes from three low-nutrient stations (11_SRF in Mediterranean Sea, 39_DCM in Arabian Sea, and 151_SRF in Atlantic Ocean) revealed expression of viral homologs of *dsr*C and *sox*YZ but not of viral P-II (Extended Data Table 1).

Overall, this systematically collected and processed GOV dataset brings critically-needed and unprecedented global ecological context to new and known, surface and deep ocean viruses. Global diversity analyses identified and mapped abundant dsDNA viruses at the population- and VC-level, and indicated that these are near-completely sampled in epipelagic waters and dominated by few viral groups, mostly newly described. The characterization of new viral-encoded AMGs, their viral carriers, possible impacted hosts, and biogeographical patterns revealed that viral manipulation of cellular processes involves much more than photosynthesis and carbon metabolism^25–28^, to also now include nitrogen and sulfur cycling throughout the epipelagic ocean. These advances are foundational for interpretation of new (meta)genomic datasets and selection of relevant experimental systems to develop, and, together with myriad experimental, informatic and theoretical advances already occurring^5,15,42–44^, will accelerate the field towards understanding and predicting the roles and global impacts of viruses in nature.

## Methods

### Sample collection and processing

#### Tara *Oceans expedition*

Ninety samples were collected between October 10, 2009, and December 12, 2011, at 45 locations throughout the world’s oceans (Supplementary Table 1) through the *Tara* Oceans Expedition^46^. These included samples from a range of depths: surface, deep chlorophyll maximum, bottom of mixed layer when no deep chlorophyll maximum was observed (Station 123, 124, and 125), and mesopelagic samples. The sampling stations were located in 7 oceans and seas, 4 different biomes and 14 Longhurst oceanographic provinces (Supplementary Table 1). For TARA station 100, two different peaks of chlorophyll were observed, so two samples were taken at the shallow (100_DCM) and deep (100_dDCM) chlorophyll maximum. For each sample, 20 L of seawater were 0.22 µm-filtered and viruses were concentrated from the filtrate using iron chloride flocculation^47^ followed by storage at 4ºC. After resuspension in ascorbic-EDTA buffer (0.1 M EDTA, 0.2 M Mg, 0.2 M ascorbic acid, pH 6.0), viral particles were concentrated using Amicon Ultra 100 kDa centrifugal devices (Millipore), treated with DNase I (100U/mL) followed by the addition of 0.1M EDTA and 0.1M EGTA to halt enzyme activity, and extracted as previously described^48^. Briefly, viral particle suspensions were treated with Wizard PCR Preps DNA Purification Resin (Promega, WI, USA) at a ratio of 0.5 mL sample to 1 mL resin, and eluted with TE buffer (10 mM Tris, pH 7.5, 1 mM EDTA) using Wizard Minicolumns. Extracted DNA was Covaris-sheared and size selected to 160–180 bp, followed by amplification and ligation per the standard Illumina protocol. Sequencing was done on a HiSeq 2000 system (101 bp, paired end reads) at the Genoscope facilities (Paris, France).

Temperature, salinity, and oxygen data were collected from each station using a CTD (Sea-Bird Electronics, Bellevue, WA, USA; SBE 911plus with Searam recorder) and dissolved oxygen sensor (Sea-Bird Electronics; SBE 43). Nutrient concentrations were determined using segmented flow analysis^49^ and included nitrite, phosphate, nitrite plus nitrate, and silica. Nutrient concentrations below the detection limit (0.02 µmol kg^−1^) are reported as 0.02 µmol kg^−1^. All data from the Tara Oceans expedition are available from ENA (for nucleotide) and from PANGAEA (for environmental, biogeochemical, taxonomic and morphological data)^50–52^.

#### Malaspina expedition

Thirteen bathypelagic samples and one mesopelagic sample were collected between April 19, 2011 and July 11, 2011 during the Malaspina 2010 global circumnavigation covering the Pacific and the North Atlantic Ocean. All samples were taken at 4,000 m depth except two samples from stations 81 and 82 collected at 3,500 and 2,150 m respectively (Supplementary Table 1). Additionally, Station M114 was sampled at the OMZ region at 294 m depth. For each sample, 80 L of seawater were 0.22 µm-filtered and viruses were concentrated from the filtrate using iron chloride flocculation^47^ followed by storage at 4°C. More details about the sampling and additional variables used in the Malaspina expedition can be found in^53^. Further processing was done as for the *Tara* Oceans samples, except that Illumina sequencing was done at DOE JGI Institute (151 bp, paired end reads).

### Dataset generation

#### Contigs assembly

An overview of the contigs generation process is provided in Supplementary Fig. 7. The first step consisted in the generation of a set of contigs using as many reads as possible from 104 oceanic viromes, including 74 epipelagic and 16 mesopelagic samples from the *Tara* Oceans expedition^6^, and 1 mesopelagic and 13 bathypelagic from the Malaspina expedition^7^. This set of contigs was generated through an iterative cross-assembly^15^ (using MOCAT^54^ and Idba_ud^55^, Supplementary Fig. 7) as follows: (i) high-quality (HQ) reads were first assembled sample by sample with the MOCAT pipeline as described in^21^, (ii) all reads not mapping (Bowtie 2^56^, options –sensitive, -X 2000, and –nondeterministic, other parameters at default) to a MOCAT contig (by which we denote ‘scaftigs’, that is, contigs that were extended and linked using the paired-end information of sequencing read^57^) were assembled sample by sample with Idba_ud (iterative k-mer assembly, with k-mer increasing from 20 to 100 by step of 20), (iii) all reads remaining unmapped to any contig were then pooled by Longhurst province (i.e. unmapped reads from samples corresponding to the same Longhurst province were gathered) and assembled with Idba_ud (with the same parameters as above), and (iv) all remaining reads unmapped from every samples were gathered for a final cross-assembly (using Idba_ud). This resulted in 10,845,515 contigs (Supplementary Fig. 7B).

#### Genome binning and re-assembly

The contigs assembled from the marine viral metagenomes might still contain redundant sequences derived from the same, or closely related populations. We set out to merge contigs derived from the same population into clusters representing population genomes. To this end, contig sequences were first clustered at 95% global average nucleotide identity (ANI) with cd-hit-est^58^(options -c 0.95 -G 1 -n 10 -mask NX, Supplementary Fig. 7B), resulting in 10,578,271 non-redundant genome fragments. Next, we used co-abundance and nucleotide usage profiles of the non-redundant contigs to further identify contigs derived from the same populations with Metabat^59^. Briefly, Metabat uses Pearson correlation between coverage profiles (determined from the mapping of HQ reads of each sample to the contigs with Bowtie 2^56^, options –sensitive, -X 2000, and –non-deterministic, other parameters at default) and tetranucleotide frequencies (Metabat parameters: 98% minimum correlation, mode “sensitive”, see Supplementary Text for more detail about the selection of these parameters) to identify contigs originating from the same genome. The 8,744 bins generated, including 3,376,683 contigs, were further analyzed, alongside 623,665 contigs not included in any genome bin but ≥1.5kb.

In an attempt to better assemble these genome bins, two additional sets of contigs were generated for each genome bin, two additional sets of contigs were generated (beyond the set of initial contigs binned by Metabat^59^), based on the de novo assembly of (i) all reads mapping to the contigs in the genome bin, and (ii) only reads from the sample displaying the highest coverage for the genome bin (both assemblies with Idba_ud^55^, Supplementary Fig. 7C). The latter might be expected to lead to the “cleanest” genome assembly because it includes the minimum between-sample sequence variation, lowering the probability of generating chimeric contig^60^. The former may be necessary if the virus is locally rare, so that sequences from multiple metagenomes are needed to achieve complete genome coverage. Thus, if the assembly from the single “highest coverage sample” was improved or equivalent (longest contig in the new assembly representing ≥95% of the longest contig in the initial assembly), this set of contigs was selected as the sequence for this bin (n=6,423). This optimal single-sample assembly was thus privileged compared to a cross-assembly (either based on the initial contigs or on the re-assembly of all sequences aligned to that bin). Otherwise, the “all samples” bin re-assembly was selected if equivalent or better than the initial assembly (longest contig representing ≥95% of the longest initial contig, n=999). The assumption that cross-assembly would be needed for locally rare viruses, without a high-coverage sample, was confirmed by the comparison between the highest coverage of these two types of bins: on average, bins for which the “optimal” assembly were selected displayed a maximum coverage of 5.47 per Gb of metagenome, while the bins for which the “cross-assembly” was selected displayed a maximum coverage of 1.37 per Gb of metagenome (Supplementary Table 2). Finally, if both re-assemblies yielded a longest contig smaller (<95%) than the one in the initial assembly, the bin was considered as a false positive (i.e. binning of contigs from multiple genomes, n=1,356), and contigs from the initial assembly were considered as “unbinned” (263,006 contigs, added to the 623,665 contigs ≥1.5kb retained as “unbinned”).

#### Identification of viral contigs and delineation of viral populations

VirSorter^61^ was used to identify microbial contigs (using the “virome decontamination” mode, with every contig ≥10kb and not identified as a viral contig being considered as a microbial contig). Sequences with a prophage predicted were manually curated to distinguish actual prophages (i.e. viral regions within a microbial contig) from contigs that belonged to a viral genome and were wrongly predicted as a prophage. Contigs originating from an eukaryotic virus were identified based on best BLAST hit affiliation of the contig predicted genes against NCBI RefseqVirus (see Supplementary Text).

The genome bins were affiliated as microbial (if 1 or more contigs were identified as microbial, n=1,763), eukaryotic virus (if contigs affiliated as eukaryotic virus comprised more than 10kb or more than 25% of the genome bin total length, n=962) or viral (i.e. archaeal and bacterial viruses, n=4,341), with the 356 remaining bins, lacking a contig long enough for an accurate affiliation, considered as “unknown”.

Viral bins were then refined to evaluate if they corresponded to a single or a mix of viral population(s). To that end, the Pearson correlation and Euclidean distance between abundance profiles

(i.e. profile of the contig average coverage depth across the 104 samples) of bin members and the bin seed (i.e. the largest contig) were computed, and a single-copy viral marker gene (TerL) was identified in binned contigs (Supplementary Fig. 7E). Thresholds were chosen to maximize the number of bins with exactly one TerL gene and minimize the number of bins with multiple TerL genes (Supplementary Fig. 7G). For each bin, contigs with a Pearson correlation coefficient to the bin seed <0.96 or a Euclidean distance to the seed >1.05 were removed from the bin, and added to the pool of unbinned contigs. Eventually, every bin still displaying multiple TerL genes after this refinement step were split, and all corresponding contigs added to the pool of “unbinned” contigs (Supplementary Fig. 7E).

The final set of contigs was formed by compiling (i) all contigs belonging to a viral bin, (ii) “unbinned” viral contigs (i.e. contigs affiliated to archaeal and bacterial virus and not part of any genome bin), and (iii) viral contigs identified in microbial or eukaryote virus bins (considered as “unbinned” contigs, Supplementary Fig. 7F). Within this set of contigs, all viral bins were considered as viral populations, as well as every unbinned viral contig ≥10kb, leading to a total of 15,222 epi- and mesopelagic populations, and 58 bathypelagic populations (Supplementary Fig. 1, Supplementary Table 2, and Supplementary Text). In this study, we focus only on the 15,222 epi- and mesopelagic populations, totaling 24,353 contigs. For the detection of AMGs, we added to these populations all short epi- and mesopelagic unbinned viral contigs (<10kb), adding up to a total of 298,383 contigs.

### Sequence clustering and annotations

#### Dataset of publicly available viral genomes and genome fragments

Genomes of viruses associated with a bacterial or archaeal host were downloaded from NCBI RefSeq (1,680 sequences, v70, 05-26-2015). To complete this dataset of reference genomes, viral genomes and genome fragments available in Genbank, but not yet in Refseq were downloaded (July 2015) and manually curated to select only bacterial and archaeal viruses (1,017 sequences). These included viral genomes not yet added to RefSeq, as well as genome fragments from fosmid libraries generated from seawater samples^12,13^. Mycophage sequences (available from http://phagesdb.org^62^) and not already in RefSeq were downloaded (July 2015) and included as well (734 sequences). Finally, 12,498 viral genome fragments from the VirSorter Curated Dataset, identified in publicly available microbial genome sequencing projects, were added to the database^9^.

#### Genome (fragments) clustering through gene-content based network analysis

Proteins predicted from 14,650 large GOV contigs (≥10kb and ≥10 genes), were added to all proteins from the publicly available viral genomes and genomes fragments gathered, and compared through all-vs-all blastp, with a threshold of 10^−5^ on e-value and 50 on bit score. Protein clusters were then defined using MCL (using default parameters for clustering of proteins, similarity scores as log-transformed e-value, and MCL inflation of 2^63^). vContact (https://bitbucket.org/MAVERICLab/ vcontact) was then used to calculate a similarity score between every pair of genome and/or contigs based on the number shared of PCs between the two sequences (as in^8,9^), and then compute a MCL clustering of the genomes/contigs based on these similarity scores (thresholds of 1 on similarity score, MCL inflation of 2). The resulting viral clusters (or VCs, clusters including ≥2 contigs and/or genomes), consistent with a clustering based on whole-genome BLAST comparison, corresponded to approximately genus-level taxonomy, with rare cases closer to subfamily-level taxonomy (Extended Data Fig. 2 and Supplementary Text). A total of 1,259 viral clusters were obtained, with 867 including at least one GOV sequence.

#### Viral contigs annotation

A functional annotation of all GOV predicted proteins was based on a comparison to the PFAM domain database (v27^64^) with HmmSearch^65^ (threshold of 30 on bit score and 1e-3 on e-value), and additional putative structural proteins were identified through a BLAST comparison to protein clusters detected in viral metaproteomics dataset^66^. This metaproteomics dataset led to the annotation of 13,547 hypothetical proteins lacking a PFAM annotation. A taxonomic annotation was performed based on a blastp of the predicted proteins against proteins from archaeal and bacterial viruses from NCBI RefSeq and Genbank (threshold of 50 on bit score and 10^−3^ on e-value).

VCs were affiliated based on isolate genome members, when available. When multiple isolates were included in the VC, the VC was affiliated to the corresponding subfamily or genus of these isolates (excluding all “unclassified” cases). This was the case for VC_2 (T4 subfamily^17,18^), and VC_9 (*T7virus*^19^). When only one or a handful of affiliated isolate genomes were included in the VC and lacked genus-level classification, a candidate name was derived from the isolate (if several isolates, from the first one isolated). This was the case for VC_5 (*Cbaphi381virus*^67^), VC_12 (*P12024virus*^68^), VC_14 (*MED4-117virus*), VC_19 (*HMO-2011virus*^69^), VC_31 (*RM378virus*^70^), VC_36 (*GBK2virus*^71^), VC_47 (*Cbaphi142virus*^67^), and VC_277 (*vB_RglS_P106Bvirus*^72^). Otherwise, VCs were considered as “new VCs”.

#### “Phage proteomic tree” (i.e. “whole-genome comparison tree”) computation and visualization

All publicly available complete genomes (see above), all complete (circular) and near-complete (extrachromosomal genome fragment >50kb with a terminase) from the VirSorter Curated Dataset, and all complete and near-complete GOV contigs were compared to generate a phage proteomic tree, as previously described^12,73^. Briefly, a proteomic similarity score was calculated for each pair of genome based on a all-vs-all tblastx similarity as the sum of bit scores of significant hits between two genomes (e-value ≤ 0.001, bit score ≥30, identity percentage ≥ 30). To normalize for different genome sizes, each genome was also compared to itself to generate a self-score, and the distance between two different genomes was calculated as a Dice coefficient (as in^12^), i.e. for two genomes A and B with a proteomic similarity score of AB, the corresponding distance d would be 1-(2*AB)/(AA+BB), with AA and BB being the self-score of genomes A and B respectively. For clarity, the tree displayed in Extended Data Fig. 2 only include non-GOV sequences found in a VC with GOV sequence(s) or within a distance <0.5 to a GOV sequence, adding for a total of 1,522 reference sequences. iTOL^74,75^ was used to visualize and display the tree.

### Distribution and relative abundance of viral populations and VCs

#### Detection and estimation of abundance for viral contigs and populations

The presence and relative abundance of a viral contig in a sample were determined based on the mapping of HQ reads to the contig sequences, computed with Bowtie 2 (options –sensitive, -X 2000, and –non-deterministic, default parameters otherwise^56^), as previously described^10^. A contig was considered as detected in a metagenome if more than 75% of its length was covered by aligned reads derived from the corresponding sample. A normalized coverage for the contig was then computed as the average contig coverage (i.e. number of nucleotides mapped to the contig divided by the contig length) normalized by the total number of bp sequenced in this sample.

The detection and relative abundance of a viral population was based on the coverage of its contigs: a population was considered as detected in a sample if more than 75% of its cumulated length was covered, and its normalized coverage was computed as the average normalized coverage of its contigs.

#### Relative abundance of VCs

The relative abundance of VCs was calculated based on the coverage of its members within the 15,222 viral populations identified. If a population included contigs all linked to the same VC, or linked to a single VC except for unclustered (because too short) contigs, this population coverage was added to the total of the corresponding VC. In the rare cases where the link between population and VC was ambiguous because different contigs within a population pointed toward different VCs (n=475, i.e. 3.1% of the populations), the population coverage was equally split between these VCs. Finally, if no contig in the population belonged to any VC (n=2,605, 17% of the populations), the population coverage was added to the “unclustered” category. Eventually, for each sample, the cumulated coverage of a VC was normalized by the total coverage of all populations to calculate a relative abundance of the VC among viral populations.

The selection of abundant VCs within a sample was based on the contribution of the VC to the sample diversity as measured by the Simpson index. For each sample, the overall Simpson index was first calculated with all VCs. Then, VCs were sorted by decreasing relative abundance and progressively added to a new calculation of the Simpson index. VCs considered as abundant were the ones which, once cumulated, represented 80% of the sample diversity (i.e. a Simpson index greater or equal to 80% of the sample total Simpson index, Extended Data Fig. 1C). The 38 VCs identified as abundant in at least 2 different stations were selected as “recurrently abundant VCs in the GOV dataset” (Fig. 2 and Extended Data Fig. 3).

### Host prediction and diversity

A genome database of putative hosts for the epi- and mesopelagic GOV viruses was generated, including all archaea and bacteria genomes annotated as “marine” from NCBI RefSeq and WGS (both times only sequences ≥5kb, 184,663 sequences from 4,452 genomes, downloaded in August 2015), and all contigs ≥5kb from the 139 *Tara* Oceans microbial metagenomes corresponding to the bacteria and archaea size fraction (791,373 sequences)^21^. For these microbial metagenomic contigs, a first blastn was computed to compare them to all GOV contigs, and exclude from the putative host dataset all metagenomic contigs with a significant similarity to a viral GOV sequence (thresholds of 50 on bit score, 0.001 on e-value, and 70% on identity percentage) on ≥90% of their length, as these are likely sequences of viral origin sequenced in the bacteria and archaea size fraction (these represented 2.2% of the contigs in the assembled microbial metagenomes). The taxonomic affiliation of NCBI genomes was taken from the NCBI taxonomy. For *Tara* Oceans contigs, a last common ancestor (LCA) affiliation was generated for each contig based on genes affiliation^21^, if 3 genes or more on the contig were affiliated. Three different approaches were then used to link viral contigs and putative host genomes (see Supplementary Text and ref. 76 for an extended discussion about the efficiency and raw results of these host prediction methods, and Supplementary Table 4 for a list of all host predictions by sequence).

#### BLAST-based identification of sequence similarity between viral contigs and host genome

All GOV viral contigs were compared to all archaeal and bacterial genomes and genome fragments with a blastn (threshold of 50 on bit score and 0.001 on e-value), to identify regions of similarity between a viral contig and a microbial genome, indicative of a prophage integration or horizontal gene transfer^76^. A host prediction was made when (i) a NCBI genomes displayed a region similar to a GOV viral contig ≥5kb at ≥70% id, or (ii) when a *Tara* Oceans microbial metagenomic contig (≥5kb) displayed a region similar to a GOV viral contig ≥2.5kb at ≥70% id.

#### Matches between GOV viral contigs and CRISPR spacers

CRISPR arrays were predicted for all putative host genomes and genome fragments (NCBI microbial genomes and *Tara* Oceans microbial metagenomic contigs) with MetaCRT^77,78^. CRISPR spacers were extracted, and all spacers with ambiguous bases or low complexity (i.e. consisting of 4 to 6 bp repeat motifs) were removed. All remaining spacers were matched to viral contigs with fuzznuc^79^, with no mismatches allowed, which although rarely observed yields highly accurate host predictions^76^.

#### Nucleotide composition similarity: comparison of tetranucleotide frequency

Bacterial and archaeal viruses tend to have a genome composition close to the genome composition of their host, a signal that can be used to predict viral-host pairs^9,76^. Here, canonical tetranucleotide frequencies were observed for all viral and host sequences using Jellyfish^80^, and mean absolute error (i.e. average of absolute differences) between tetranucleotide frequency vectors were computed with in-house Perl and Python scripts for each pair of viral and host sequence. A GOV viral contig was then assigned to the closest sequence (i.e. lowest distance d) from the pool of NCBI genomes if d<0.001 (because both the tetranucleotide frequency signal and the taxonomic affiliation of these complete genomes are more robust than for metagenomic contigs), and otherwise assigned to the closest (i.e. lowest distance) *Tara* Oceans microbial contig if d<0.001.

#### Summarizing host prediction at the VC level

Overall, 3,675 GOV contigs could be linked to a putative host group among the 24,353 GOV contigs associated with an epi- or mesopelagic viral population. To summarize these affiliations at the VC level, a Poisson distribution was used to estimate the number of expected false positive associations for each VC – host group combination based on (i) the global probability of obtaining a host prediction across all pairs of viral and host sequences tested and for all methods (p=5.8x10^−08^), (ii) the number of potential predictions generated for the VC, corresponding to 3 times the number of sequences in the VC (to take into account the three methods), and (iii) the number of sequences from the host group in the database. By comparing the number of links observed between a VC and a host group to this expected value, which takes into account the bias in database (i.e. some host groups will be over- or under-represented in our set of archaeal and bacterial genomes and genome fragments) and the bias linked to the variable number of sequences in VCs, we can determine if the number of associations observed for any VC – host group combination is likely to be due to chance alone (and calculate the associated p-value).

#### Microbial community diversity and richness indexes

Diversity and richness indexes for putative host populations were based on the OTU abundance matrix generated from the analysis of _mi_TAGs in *Tara* Oceans microbial metagenomes^21^. These indexes were computed for each host group at the same taxonomic level as the host prediction, i.e. the phylum level except for Proteobacteria where the class level is used. The R package vegan^81^ was used to estimate for each group (i) a global Chao index (i.e. including all OTUs from all samples) through the function estaccumR, (ii) a sample-by-sample Chao index with the function estimateR, and (iii) Sorensen indexes between all pairs of samples with the function betadiver. Diversity indexes presented in Extended Data Fig 4 were based on epipelagic samples only, as the 38 VCs identified as abundant were mostly retrieved in epipelagic samples. Candidate division OP1 was excluded from this analysis because no OTU affiliated to this phylum was identified.

### Identification and annotation of putative AMGs

#### Detection of AMGs

Predicted proteins from all GOV viral contigs were compared to the PFAM domain database (hmmsearch^65^, threshold of 40 on bit score and 0.001 on e-value), and all PFAM domains detected were classified into 8 categories: “structural”, “DNA replication, recombination, repair, nucleotide metabolism”, “transcription, translation, protein synthesis”, “lysis”, “membrane transport, membrane-associated”, “metabolism”, “other”, and “unknown” (as in ^24^). Four AMGs (i.e. similar to a domain from the “metabolism” category) were then selected for further study because of their central role in sulfur (*dsr*C and *sox*YZ) or nitrogen (P-II, *amo*C) cycle, and the fact that these had never been detected in a surface ocean viral genome so far (*dsrC/tusE*-like genes have been detected in deep water viruses^14,29^). To evaluate if an AMG was “known”, a list of PFAM domain detected in NCBI RefSeqVirus and Environmental Phages was computed based on a similar hmmsearch comparison (threshold of 40 on bit score and 0.001 on e-value), and augmented by manual annotation of AMGs from^24,82^. The complete list of PFAM domains detected in GOV viral contigs is available as Supplementary Table 5.

#### Phylogenetic tree generation and contigs map comparison

Sequences similar to these AMGs were recruited from the *Tara* Oceans microbial metagenomes^21^ based on a blastp of all predicted proteins from microbial metagenome to the viral AMGs identified (threshold of 100 on bit score, 10^−5^ on e-value, except for P-II where a threshold of 170 on bit score was used because of the high number of sequences recruited). The viral AMG sequences were also compared to NCBI nr database (blastp, threshold of 50 on bit score and 10^−3^ on e-value) to recruit relevant reference sequences (up to 20 for each viral AMG sequence). These sets of viral AMGs and related protein sequences were then aligned with Muscle^83^, the alignment manually curated to remove poorly aligned positions with Jalview^84^, and two trees were computed from the same curated alignment: a maximum-likelihood tree with FastTree (v2.7.1, model WAG, other parameters set to default^85^) and a bayesian tree with MrBayes (v3.2.5, mixed evolution models, other parameters set to default, 2 MCMC chains were run until the average standard deviation of split frequencies was <0.015, relative burn-in of 25% used to generate the consensus tree^86^). In all cases except AmoC, the mixed model used by MrBayes was 100% WAG, confirming that this model was well suited for archaeal and bacterial virus protein trees. Manual inspection revealed only minor differences between each pair of trees, so an SH test was used to determine which tree best fitted the sequence alignment, using the R library phangorn^87^. Itol^74^ was used to visualize and display these trees, in which branches with supports <40% were collapsed. Annotated interactive trees are available online at http://itol.embl.de/shared/Siroux. Contigs map comparison were generated with Easyfig^88^, following the same method as for the VCs (see Supplementary Information).

#### Functional characterization of putative AMGs

Conserved motifs were identified on the different AMGs based on the literature: *dsr*C conserved motifs were obtained from ref. 33, *sox*YZ conserved residues were identified from the PFAM domains PF13501 and PF08770, and P-II conserved motifs from PROSITE documentation PDOC00439. A 3D structure could also be predicted for P-II AMGs by I-TASSER^89^ (default parameters), the quality of these predictions being confirmed with ProSA web server^90^. To further confirm the functionality of these genes, selective constraint on these AMGs was evaluated through pN/pS calculation, as in ref. 91. Briefly, synonymous and non-synonymous SNPs were observed in each AMG, and compared to expected ratio of synonymous and non-synonymous SNPs under a neutral evolution model for this genes. The interpretation of pN/pS is similar as for dN/dS analyses, with the operation of purifying selection leading to pN/pS values < 1. Finally, AMG transcripts were searched in metatranscriptomic datasets generated through the *Tara* Oceans consortium (ENA Id ERS1092158, ERS488920, and ERS494518). Briefly, For 0.2–1.6 and 0.22–3µm filters, bacterial rRNA depletion was carried out on 240–500 ng total RNA using Ribo-Zero Magnetic Kit for Bacteria (Epicentre, Madison, WI). The Ribo-Zero depletion protocol was modified to be adapted to low RNA input amounts^92^. Depleted RNA was used to synthetize cDNA with SMARTer Stranded RNA-Seq Kit (Clontech, Mountain View, CA)^92^. Metatranscriptomic libraries were quantified by qPCR using the KAPA Library Quantification Kit for Illumina Libraries (KapaBiosystems, Wilmington, MA) and library profiles were assessed using the DNA High Sensitivity LabChip kit on an Agilent Bioanalyzer (Agilent Technologies, Santa Clara, CA). Libraries were sequenced on Illumina HiSeq2000 instrument (Illumina, San Diego,CA) using 100 base-length read chemistry in a paired-end mode. High quality reads were then mapped to viral contigs containing *dsrC, soxYZ,* P-II, or *amo*C genes with SOAPdenovo2^57^ within MOCAT^54^ (options *screen* and *filter* with length and identity cutoffs of 45 and 95%, respectively, and paired-end filtering set to *yes*), and coverage was defined for each gene as the number of bp mapped divided by gene length (including only reads mapped to the predicted coding strand).

#### Distribution of AMGs and association with geochemical metadata

The distribution and relative abundance of AMGs was based on the read mapping and normalized coverage of the contig including the AMG. To get a range of temperature and nutrient concentrations for the widespread AMGs (detected in >5 stations) that takes into account both the samples in which these AMGs were detected and the differences in normalized coverage, a set of samples was selected through a weighted random drawing replacement, with the weight of each sample corresponding to the AMG’s normalized coverage. That way, a range of temperature or nutrient concentration values associated with the AMG’s distribution and abundance could be generated for each AMG and each environmental parameter tested. The number of samples randomly selected for each AMG was the same as the total number of samples for which a value of this parameter was available.

### Scripts and data availability

Scripts used in this manuscript are available on the Sullivan lab bitbucket under project “GOV_Ecogenomics” (https://bitbucket.org/MAVERICLab/gov_ecogenomics). All raw reads are available through ENA (*Tara* Oceans) or JGI (Malaspina) using the dataset Ids listed in Supplementary Table 1. Processed data are available through iVirus, including all sequences from assembled contigs, list of viral populations and associated annotated sequences as genbank files, viral clusters composition and characteristics, map comparisons of genomes and contigs of the 38 abundant VCs, and host predictions for viral contigs.

## Extended Data

**Extended Data Figure 1: Accumulation curves of populations (A) and viral clusters (VCs, B) and identification of abundant VCs in GOV samples (C).** A & B. Accumulation curves were computed from 50 random shuffling of samples (blue dots), either with all, epipelagic, mesopelagic, or bathypelagic samples. For each curve, the average of 50 iterations is highlighted with red dots. C. Schematic of the selection process of abundant VCs. For each sample, VCs accounting for (up to) 80% of the sample diversity (as assessed by Simpson index) were considered as abundant (example for sample 125_MIX on the left). VCs detected as abundant in at least two different stations were included in the 38 VCs described in Fig. 2 and Extended Data Fig. 3.

**Extended Data Figure 2: Comparison of VCs with other classification methods: phage proteomic tree and percentage of shared genes.** The phage proteomic tree includes the 756 GOV complete and near-complete genomes from epi- and mesopelagic samples, and closest references from RefSeq and Environmental phages (d<0.5 to a GOV sequence or found in the same VC as a GOV sequence). All VCs with more than 8 representatives in the tree or part of the 38 abundant VCs are indicated with coloring of the outer ring. The name and affiliation (if available) of the 38 abundant VCs are indicated next to the VC on the colored ring. Branches of monophyletic clades including more than 3 GOV and/or uncultivated marine sequences with no isolate reference are highlighted in blue. Inset: distribution of number of shared genes (i.e. number of shared protein clusters) for viral genome/contigs pairs either between different VCs or within VCs.

**Extended Data Figure 3: Summary of 34 of the 38 abundant viral clusters (VCs, the 4 other abundant VCs being the ubiquitous ones presented in Fig. 2).** Predicted genome size is based on the set of isolates and circular contigs in the VC (NA corresponds to VCs without any circular contigs, or for which the relative standard deviation of estimated genome size across the different isolate(s) and/or circular contigs is greater than 15%). Host association values are based on the number of cluster members associated with each host group, the statistical significance of this number of predictions being evaluated by comparison with an expected number of associations calculated from a Poisson distribution. Host associations based on known isolates are indicated with a star (for associations based on cultivated isolates) or a dot (for associations based on the detection of a cluster member in a microbial genome from the VirSorter Curated Dataset). The abundant epipelagic microbial groups (representing >1% of the microbial OTUs abundance of epipelagic samples) are highlighted in bold. Distribution and relative abundance of VCs are based on the cumulated coverage of VC members among sample viral populations. The main oceanic basins are indicated for each set of sample, Med. Sea-Mediterranean Sea.

**Extended Data Figure 4: Association between abundant viral clusters (VCs) and host group abundance and diversity A.** Abundance and diversity of bacterial and archaeal host groups associated with the 38 abundant VCs (see Fig. 2A). For each host group (phylum level, except for Proteobacteria where the class level is used), the different panels display from top to bottom (i) the number of VCs associated with this host group, (ii) the global relative abundance of this group estimated from the microbial metagenomic OTU counts, (iii) the global diversity of this group based on a Chao index computation including all *Tara* Oceans microbial metagenome samples (i.e. including both Alpha and Beta diversity), (iv) the distribution of Chao indexes by sample for this group (Alpha diversity), and (v) the average Sorensen index between pairs of samples including at least one OTU of this group (Beta diversity). OTU counts were derived from the 109 epipelagic microbial metagenomes described in^21^. **B.** Pearson correlations between host group relative abundance or diversity indexes (Global Chao, Average Chao across samples, and Average Sorensen across samples) and the number of VCs.

**Extended Data Figure 5: Diversity, distribution, and genome context of *dsr*C genes in GOV contigs. A.** Maximum-likelihood tree (from an amino-acid alignment) including the 11 viral DsrC and microbial sequences from microbial metagenomes and NCBI nr database. The presence of conserved C residues (named Cys-A & Cys-B, as in^33^) is indicated with color circles next to each sequence or clade, and the corresponding type of DsrC-like protein is indicated by coloring the branch or clade. The microbial metagenomic contigs affiliated to uncultivated, marine sulfur-oxidizing Gammaproteobacteria (as confirmed by complementary phylogenetic analysis of DsrAB, Supplementary Fig. 6) are indicated with a star next to the sequence or clade. Viral AMG sequences are highlighted in blue, internal nodes SH-like supports are represented by proportional circles (all nodes with support < 0.40 were collapsed). Each *dsrC* AMG is associated with an abundance profile (on the right) displaying the relative abundance of the contig across the 91 epi- and mesopelagic samples (based on normalized coverage, i.e. contig coverage / Gb of metagenome). **B.** Comparison of *dsrC-*containing contigs maps. T4-like marker genes (PhoH and T4 baseplate) are indicated on the maps, alongside putative AMGs (Fe-S biosyn for Iron-sulfur cluster biosynthesis, and Amt for Ammonia transporter).

**Extended Data Figure 6: Diversity, distribution, and genome context of s*ox*YZ genes in GOV contigs. A.** Bayesian tree (from an amino-acid alignment) including the 4 viral SoxYZ and microbial sequences from microbial metagenomes and NCBI nr database. The affiliation of microbial clades (either from the NCBI reference or from the LCA affiliation of metagenomic contigs) is indicated by coloring of the grouped clades or with a colored square next to the sequence. Viral AMG sequences are highlighted in blue, posterior probabilities are represented by proportional circles (all nodes with posterior probability < 0.40 were collapsed). Clades including sulfur-oxidation proteobacteria are indicated on the tree. Each s*ox*YZ AMG is associated with an abundance profile (on the right) displaying the relative abundance of the contig across the 91 epi- and mesopelagic samples (based on normalized coverage, i.e. contig coverage / Gb of metagenome). **B.** Comparison of *sox*YZ-containing contigs maps. For contig GOV_bin_4310_contig-100_0, the second largest contig from the same bin (GOV_bin_4310_contig-100_1) is displayed. T4-like marker genes (PhoH, Gp23 and T4 baseplate) are indicated on the maps, alongside putative AMGs (Fe-S biosyn: Iron-sulfur cluster biosynthesis).

**Extended Data Figure 7: Diversity, distribution, and genome context of P-II genes in GOV contigs. A.** Maximum-likelihood tree (from an amino-acid alignment) including the 10 viral P-II and microbial sequences from microbial metagenomes and NCBI nr database. The affiliation of microbial clades (either from the NCBI reference or from the LCA affiliation of metagenomic contigs) is indicated by coloring of the grouped clades or with a colored square next to the sequence. The sequences lacking the conserved uridylation site of P-II (Supplementary Fig. 4) are highlighted with a star next to the sequence name or clade. Viral AMG sequences are highlighted in blue, internal nodes SH-like supports are represented by proportional circles (all nodes with support < 0.40 were collapsed). Each P-II AMG is associated with an abundance profile (on the right) displaying the relative abundance of the contig across the 91 epi- and mesopelagic samples (based on normalized coverage, i.e. contig coverage / Gb of metagenome). **B.** Comparison of P-II-containing contigs maps. Ammonia transporter genes linked to P-II are indicated on the map (Amm Transp, dark red). When available, the VC affiliation of each contig is indicated next to the contig name. Contig GOV_bin_5834_contig-100_7 is too short to be clustered based on a shared PC network, however the seed contig of its population was clustered (in VC_12, *Siphoviridae - P12024virus*), hence this seed contig affiliation is indicated.

**Extended Data Figure 8: Diversity, distribution, and genome context of *amo*C gene in GOV contigs. A.** Maximum-likelihood tree (from an amino-acid alignment) including the GOV *amo*C AMG and microbial sequences from microbial metagenomes and NCBI nr database. The affiliation of microbial clades (either from the NCBI reference or from the LCA affiliation of metagenomic contigs) is indicated by coloring of the grouped clades or with a colored square next to the sequence. Viral AMG sequence is highlighted in blue, internal nodes SH-like supports are represented by proportional circles (all nodes with support < 0.40 were collapsed). **B.** Abundance profile displaying the relative abundance of the contig across the 91 epi- and mesopelagic samples (based on normalized coverage, i.e. contig coverage / Gb of metagenome). **C.** Map of the *amo*C-containing contig.

**Extended Data Figure 9: Normalized coverage of contigs harboring AMG as function of the temperature and nutrient concentrations (NO_2_, NO_3_, PO_4_) of the corresponding samples.** AMGs are grouped by clade based on the phylogeny (see Extended Data Fig. 5-6-7), and coverages are cumulated when a clade included multiple contigs. Plots display the cumulated normalized coverage of a clade (y-axis) as function of the temperature or nutrient concentration (x-axis) across all epipelagic samples (mesopelagic samples were excluded from the analysis since the AMG signal was detected in epipelagic samples), only for clades not geographically restricted (i.e. found in >5 samples, see Fig. 3C). Samples are color-coded according to their ocean and sea region (Supplementary Table 1). The calculated preferential range of temperature or nutrient concentration is displayed below each plot for the epipelagic AMGs (P-II-4 distribution could not be linked to specific environmental conditions, but this AMG is the only one consistently retrieved in mesopelagic samples).

**Extended Data Table 1: Summary of genes and contigs characteristics for new viral DsrC, SoxYZ, and P-II AMGs.** Each gene is linked to its contig, and when available, to the corresponding viral population and predicted host (from BLAST hit, CRISPR spacer similarity, or nucleotide composition similarity). Widespread and abundant VCs are highlighted in bold. In addition, the calculated pN/pS of each gene is indicated (measuring the strength of selection pressure occurring for this gene, the gene with a pN/pS not representing a strong purifying selection is highlighted in red), as well as the coverage of these genes and other genes in the contigs in 3 metatranscriptomic samples from 3 open ocean Tara stations (cases where the AMG coverage is >0.5 and associated with the coverage of other genes from the same viral contig are highlighted in green).

## Supplementary Information

### Supplementary Text

**Supplementary Table 1: List of viromes included in the GOV dataset.** For each virome, the corresponding expedition, station number, and depth is indicated. *Tara* Oceans stations are prefixed with “Tara_” and Malaspina stations with an “M”. Accession numbers are given for raw reads available in ENA (for *Tara* Oceans samples) and on JGI IMG (for *Malaspina* samples). Longhurst provinces and biomes are defined based on Longhurst^93^ and environmental features are defined based on Environment Ontology (http://environmentontology.org/). The total number of reads and bp sequenced, as well as the number of bp mapped to viral contigs within and outside of populations are indicated. *Malaspina stations for which no water mass or basin data are available because these were not included in the previous study^53^.

**Supplementary Table 2: GOV viral population summary.** The number of contigs, total length and length of the largest contig, type of assembly used, and highest normalized coverage across the GOV samples is indicated in the first tab. For populations already identified in the TOV dataset (contigs similar at 95% ANI on ≥50% of their length), the size of the TOV contig is noted. In the second tab, the normalized coverage (average coverage of the population contig(s) normalized by the total sequencing depth of the sample) is indicated as coverage / Gb of metagenome for all GOV samples.

**Supplementary Table 3: Summary of Viral Clusters (VCs).** The first tab lists, for each VC, the number of members (total, and by dataset, i.e. originating from RefSeq, environmental phages, VirSorter Curated Dataset, and GOV), alongside the affiliation of RefSeq members of the VCs (when available) at the family, subfamily, and genus levels. The second tab includes the cumulative normalized coverage of each VC in each sample (based on the coverage of populations members of the VC), as well as the sum of coverage for the 38 recurrently abundant VCs and all other VCs at the bottom.

**Supplementary Table 4: List of host prediction for GOV viral contigs associated with a population.** For each prediction, the type of signal (blastn, CRISPR, tetranucleotide composition), the host sequence used for the prediction alongside its affiliation, and the strength of the prediction (length of the blastn match, number of mismatches in the CRISPR spacer, and distance between viral and host tetranucleotide frequencies vectors) is indicated.

**Supplementary Table 5: List of PFAM domains detected in GOV viral contigs.** For each PFAM domain, the number of genes detected in the GOV dataset is indicated, alongside the category of the domain (as in ^24^). The category “other” category includes PFAM domains with vague descriptions, multiple functions, or regulatory functions.

**Supplementary Figure 1: Schematic of the different levels of organization used in this study.** The base unit is the contig, i.e. assembled genome (fragment). These contigs are gathered (when available) in viral populations, a proxy for viral “species”, through genome binning based on co-occurrence and similarity in nucleotide composition. A higher level of organization (VCs, subfamily ~ genus level) is achieved by clustering the contigs based on shared gene content.

**Supplementary Figure 2: Multiple alignment of *dsr*C protein sequences.** Conserved residues are indicated below the alignment, and the two conserved C residues representing the active sites of “true *dsr*C” (Cys-B and Cys-A) are named as in^33^. Viral AMGs are highlighted in bold, with previously described anoxic SUP05 viruses sequences in red (from^14,29^) and epipelagic GOV sequences in black.

**Supplementary Figure 3: Multiple alignment of *soxYZ* protein sequences.** Conserved residues are indicated below the alignment for SoxY and SoxZ protein domains, based on the respective PFAM domains (PF13501 and PF08770). Viral AMGs are highlighted in bold.

**Supplementary Figure 4: Alignment (A) and predicted 3D structures (B) of P-II AMGs.** Conserved motifs are indicated below the alignment (PROSITE: PDOC00439). The uridylation site is highlighted with a star. Characterized structure (from E. Coli) and predicted 3D conformations are colored according to secondary structures (alpha helix: blue, beta strand: red), except for the trimer structure of E. Coli PII where each subunit is colored differently. For predicted structures, the model quality as assessed by ProSA^90^ is indicated below the model. Viral AMGs are highlighted in bold.

**Supplementary Figure 5: Alignment (A) and predicted transmembrane domain (B) of *amo*C AMGs.** The viral sequence is highlighted in bold, and conserved residues are indicated below the alignment. Transmembrane domains were predicted with TMHMM^94^ for the AMG *amo*C (left), and a reference *amo*C from the ammonia-oxidizing *Nitrosopumilus maritimus* SCM1 (right).

**Supplementary Figure 6: Dissimilatory sulfite reductase (*dsrAB*) tree showing the phylogeny of oxidative bacterial type *dsrAB***. Sequences from Tara Ocean microbial metagenomes close to *dsrC-*5 AMG are colored in blue and are affiliated with sulfur-oxidizing Gammaproteobacteria. Other phylogenetic groups and *dsrAB* families are collapsed and shown as triangles.

**Supplementary Figure 7: Overview and result of the cross assembly, binning, and viral contigs selection process. A.** Iterative assembly viromes. First, for each sample, reads were mapped to the set of contig generated through MOCAT^54^. Reads not assembled (i.e. not mapped to any contigs) were then used in another assembly, using Idba_ud^55^. Unmapped reads after this second round of sample-by-sample assembly were then pooled by Longhurst province (i.e. all unmapped reads from all samples within one province), and cross-assembled with Idba_ud^15^. Finally, all unmapped reads after this third round of assembly were gathered and assembled with Idba_ud. **B.** Results of the iterative assembly process. For each assembly round, the number of contigs is displayed alongside the cumulated percentage of reads mapped to a contig. **C.** Overview of the binning process. Contigs generated through the iterative assembly were binned based on correlation between their abundance profile and similarities between their tetranucleotide frequency (using Metabat^59^). For each bin, two contig pools (beyond the initial set of contigs) were generated, assembling either all reads mapping to the contig pool, or only reads from the sample in which the bin had the highest coverage (both assemblies computed with Idba_ud). The set of contigs including the largest genome fragment was then kept for each bin. **D.** Results of the re-assembly of bins. For each type of bin assembly (highest coverage sample, all samples, or initial assembly) the number of bins for which this type was selected is indicated on top, with the distribution of increase in length of longest contig at the bottom. **E.** Bin refinement based on abundance profile similarities. For each bin, the abundance profile of each contig was compared to the abundance profile of the bin seed contig (largest contig), and contigs not well correlated to the bin seed were excluded. Bins still displaying multiple TerL gene (single-copy marker gene for viruses) after this bin refinement step were split. **F.** Bin affiliation and viral population definition. Bins were either affiliated as entirely viral and considered as single viral populations, or included non-viral contigs, in which case viral contigs in these bins were considered as “unbinned” and selected as viral population seed if ≥10kb **G.** Selection of thresholds for bin refinement based on abundance profile similarities. Thresholds to exclude contigs from bins based on Euclidean distance and Pearson correlation coefficient between contig abundance profile and bin seed profile were explored, looking for the best compromise between number of true positive (z-axis, number of bins with a single TerL) and number of false negative (in colors, number of bins with multiple TerL). The thresholds combination chosen is indicated with a black square.

## Acknowledgements

We thank Joshua Weitz for advice on statistics, Claus Pelikan for help with the DsrAB phylogenetic tree, Christiane Dahl for valuable discussion regarding DsrC function, and members of the Sullivan and Rich labs for suggestions and comments on this manuscript. We acknowledge support from UA high-performance computing and the Ohio Supercomputer Center. This viral research was funded by a National Science Foundation grant (#1536989) and Gordon and Betty Moore Foundation grants (#3790, #GBMF2631) to MBS. SR was partially supported by the University of Arizona Technology and Research Initiative Fund through the Water, Environmental and Energy Solutions Initiative and the Ecosystem Genomics Institute. BED was supported by the Netherlands Organization for Scientific Research (NWO) Vidi grant 864.14.004 and CAPES/BRASIL. AL was supported by the Austrian Science Fund (FWF, project P25111-B22). We thank all people who prepared virus samples on board: Silvia G Acinas, Defne Arslan, Lucie Bittner, Julia Boras, Encarna Borruell, Christophe Boutte, Camille Clerissi, Montserrat Coll, Francisco M. Cornejo-Castillo, Cristina Díez Vives, Celine Dimier, Melissa Beth Duhaime, Isabel Ferrera, Laurence Garczarek, Nigel Grimsley, Pascal Hingamp, Lee Karp-Boss, Elena Lara, Noan LeBescot, Thomas Lefort, Hiroyuki Ogata, Eric Pelletier, Julie Poulain, Frank Prejger, Guillem Salazar, Lucie Subirana, Anne Thompson, Dolors Vaqué, Emilie Villar, as well as Montse Sala for organizing the transfer and shipping of Malaspina samples. The *Tara* Oceans consortium effort was possible thanks to the commitment of the following people and sponsors: CNRS (in particular Groupement de Recherche GDR3280), European Molecular Biology Laboratory (EMBL), Genoscope/CEA, VIB, Stazione Zoologica Anton Dohrn, UNIMIB, Fund for Scientific Research – Flanders, Rega Institute, KU Leuven, The French Ministry of Research, the French Government ‘Investissements d’Avenir’ programmes OCEANOMICS (ANR-11-BTBR-0008), FRANCE GENOMIQUE (ANR-10-INBS-09-08), MEMO LIFE (ANR-10-LABX-54), PSL* Research University (ANR-11-IDEX-0001-02), ANR (projects POSEIDON/ANR-09-BLAN-0348, PROMETHEUS/ANR-09-PCS-GENM-217, TARA-GIRUS/ANR-09-PCS-GENM-218), European Union FP7 (MicroB3/No.287589), ERC Advanced Grant Award to CB (Diatomite: 294823), BIO5, and Biosphere 2 to MBS, Spanish Ministry of Science and Innovation grant CGL2011-26848/BOS and MicroOcean PANGENOMICS to SGA, TANIT (CONES 2010-0036) from the Agència de Gestió d ‘Ajuts Universitaris i Recerca to SGA, JSPS KAKENHI Grant Number 26430184 to HO. We also thank the support and commitment of Agnès b. and Etienne Bourgois, the Veolia Environment Foundation, Region Bretagne, Lorient Agglomeration, World Courier, Illumina, the EDF Foundation, FRB, the Prince Albert II de Monaco Foundation, the *Tara* schooner and its captains and crew. We thank MERCATOR-CORIOLIS and ACRI-ST for providing daily satellite data during the expedition. We are also grateful to the French Ministry of Foreign Affairs for supporting the expedition and to the countries who graciously granted sampling permissions. *Tara* Oceans would not exist without continuous support from 23 institutes (http://oceans.taraexpeditions.org). We also acknowledge excellent assistance from the European Bioinformatics Institute (EBI), as well as Yan Yuan and the EMBL IT core facility. Malaspina Expedition and Malaspinomics Projects are funded by the Spanish Ministry of Economy and Competitiveness (refs. CSD2008-00077 and CTM2011-15461-E) to CMD. Sequencing of the Malaspina viruses metagenomes was done at the Joint Genome Institute and was supported by US Department of Energy Joint Genome Institute 2011 Microbes Program grant CSP 602 to SGA. The work conducted by the US Department of Energy Joint Genome Institute is supported by the Office of Science of the US Department of Energy under Contract No. DE-AC02–05CH11231. The authors further declare that all data reported herein are fully and freely available from the date of publication, with no restrictions, and that all of the samples, analyses, publications, and ownership of data are free from legal entanglement or restriction of any sort by the various nations whose waters the *Tara* Oceans expedition sampled in. Data described herein is available at EBI and PANGAEA (see Supplementary Table 1). All authors approved the final manuscript. This article is contribution number XXX of the *Tara* Oceans Expedition.

## Author Contributions

S.R., and M.B.S. designed the study. C.D., M.P., and Sa.S., contributed extensively to sampling collection. S.K-L. managed the logistic of the *Tara* Oceans project. B.T.P., N.S. and E.L. performed the viral-specific processing of the samples. J.P., C.C., A.A., and P.W. lead the sequencing of viral samples. S.R., S.S. and B.E.D. lead the assembly of raw data. S.R., S.S., M.B.D. and M.B.S. analyzed the genomic diversity data. S.R., A.L., J.R.B. and M.B.S. analyzed the AMGs data. S.R., J.R.B., B.E.D, S.S., M.B.D., A.L., S.P., P.B., S.G.A., C.D., J.M.G., D.V. and M.B.S. provided constructive comments, revised and edited the manuscript. *Tara* Oceans coordinators provided creative environment and constructive criticism throughout the study. All authors discussed the results and commented on the manuscript.

## Author Information

All raw data are available at ENA or IMG, with sample identifiers indicated in Supplementary Table 1. Processed data, including assembled contigs, populations definition and abundance, clusters definition and abundance, and all annotated viral contigs are available at iVirus, http://mirrors.iplantcollaborative.org/browse/iplant/home/shared/ivirus. All scripts are available at MAVERICLab bitbucket: http://bitbucket.org/MAVERICLab/gov_ecogenomics/overview. Reprints and permissions information is available at www.nature.com/reprints. Correspondence and requests for materials should be addressed to.

